# Comparison of visual quantities in untrained deep neural networks

**DOI:** 10.1101/2022.09.08.507097

**Authors:** Hyeonsu Lee, Woochul Choi, Dongil Lee, Se-Bum Paik

## Abstract

The ability to compare quantities of visual objects with two distinct measures, proportion and difference, is observed in newborn animals. Nevertheless, how this function originates in the brain, even before training, remains unknown. Here, we show that neuronal tuning for quantity comparison can arise spontaneously in completely untrained deep neural networks. Using a biologically inspired model neural network, we found that units selective to proportions and differences between visual quantities emerge in randomly initialized networks and that they enable the network to perform quantity comparison tasks. Further analysis shows that two distinct tunings to proportion and difference both originate from a random summation of monotonic, nonlinear responses to changes in relative quantities. Notably, we found that a slight difference in the nonlinearity profile determines the type of measure. Our results suggest that visual quantity comparisons are primitive types of functions that can emerge spontaneously in random feedforward networks.

**One sentence summary:** The ability to compare visual quantities arises spontaneously in untrained deep neural networks.

**Research Highlights:**

- The ability to compare visual quantity arises spontaneously in untrained networks
- Distinct tunings to measure proportion and difference of quantities are observed
- Random wiring of monotonic, nonlinear activity induces quantity-comparison units
- The nonlinearity pattern of the source unit determines the type of target measure

## Introduction

Quantity comparison, the ability to compare numbers of visual objects in two or more groups, is considered to be an essential cognitive function for the survival of animals as they hunt and forage^1–8^. For example, a group of herd animals such as lions or chimpanzees make decisions as to whether or not to approach their opponents based on the relative size of the groups^9–11^. Such behavior requires arithmetic skills to compare discrete ratios, distinct from simple magnitude perception^12–15^. Specifically, it has been observed that many different animals, including humans, can detect the ratio between two discrete quantities as well as their absolute difference^16–25^; certain animals use proportional information to guide their behavior, as is well characterized by the distance effect and Weber-Fechner law^8,9^. These reports suggest that quantity comparisons in the brain can be executed by neural activities that measure the ratio and difference between two quantities.

Subsequent studies using single-unit recordings in rhesus monkeys and fMRI of the human parietal lobe found neurons that selectively respond to a certain proportion between two non-symbolic visual quantities^18,26,27^. These neurons’ responses were observed to be largely activated when the monkeys correctly compared two proportional quantities^26,27^, implying that these proportion-selective neurons play crucial roles in visual quantity comparisons. Importantly, several studies reported that even newborn animals and human infants can perceive both absolute differences and ratios between two quantities^28–31^, suggesting that neural selectivity for quantity comparison emerges even before training or visual experience. However, it remains not fully understood how these capabilities arise initially.

In general, studying the origin of innate brain functions is challenging. Particularly, finding novel functions such as proportion- and difference-selective neural tunings in young animal brains requires extensive work that includes delicate experiments. Moreover, the corresponding developmental mechanisms can also be extremely elusive. Under such conditions, a computational model approach can provide an effective solution^32,33^. A recent study suggested the use of AlexNet^34^, a widely used convolutional network model designed following the ventral visual pathway of the brain, to explore the developmental mechanisms of innate visual functions^35,36^. In particular, one of these studies^35^ reported that neuronal tuning to visual quantities can emerge in an untrained AlexNet without any type of training. The following analysis demonstrated that abstract number sense can arise spontaneously from a random architecture of hierarchical feedforward networks, an outcome not understandable without the help of results from computational model simulations.

By adopting a similar approach, i.e., a computational model simulation, here we study whether quantity comparison functions can emerge spontaneously without training. We show that neuronal units that selectively respond to proportions and differences between two non-symbolic visual quantities emerge in a randomly initialized AlexNet in the complete absence of training. We found that two types of units respond robustly to preferred proportion and difference values regardless of the total number of stimulus objects or variations of low-level visual features, such as the size of the objects. These selective units enable the network to perform a visual quantity comparison task while also reproducing the behavioral and neuronal characteristics observed in animal experiments^26,27^.

To understand how these units can emerge in the absence of learning, we introduce a theoretical model of the spontaneous emergence of neural tuning demonstrating that quantity comparison functions can emerge from a projection of monotonically increasing and decreasing neural activities that arise spontaneously from the statistical bias of random feedforward wiring in deep hierarchical networks. We validated this scenario by confirming that the observed comparison units indeed receive strong inputs from increasing and decreasing units in the previous layer and that ablating these connections suppresses emergence of comparison units. Importantly, we found that a slightly different pattern of these increasing and decreasing neural activities leads to emergence of two distinct types of selectivity: proportion and difference tuning.

Taken together, our results suggest that visual quantity comparison ability can arise spontaneously in untrained deep neural networks and that various types of primitive functions originate from bottom-up projections in a random hierarchical network.

## Results

### Model neural networks and controlled stimuli for visual quantity comparisons

We used computational model approaches to answer the following questions: Can quantity comparison tunings, such as proportion- and difference-selective neural activities, emerge in untrained networks in the complete absence of learning? If so, what is the developmental mechanism underlying their spontaneous emergence? To answer these questions, we used a model simulation of a biologically inspired deep neural network (DNN) designed following the ventral visual pathway of the brain^34–37^. To observe two distinct types of tunings, proportion and difference selectivity, we designed a set of non-symbolic numerical stimuli composed of white and black dots (N_dots_ = N_white_ + N_black_ = 6, 12, or 18) in random spatial positions (Fig. 1a). We prepared seven different sets with distinct values of proportions (P = N_white_ / N_dots_ = 0/6, 1/6, 2/6, 3/6, 4/6, 5/6, 6/6) and differences (D = N_white_ - N_black_ = −6, −4, −2, 0, +2, +4, +6), with each condition containing 200 images of different dot distributions. Multiple sets of images for different total numbers of dots were prepared to examine neural responses separately for each proportion and difference value (e.g., for images with (N_white_, N_black_) = (1, 5) and (2, 10), P is 1/6 in both cases, but D differs at −4 and −8). As a model of young brain networks before sensory experience, we used a randomly initialized untrained AlexNet in which the connectivity weights in all layers were randomly selected^35,36^, meaning that convolutional filters do not form any particular shape (Fig. 1b).

**Figure 1.**
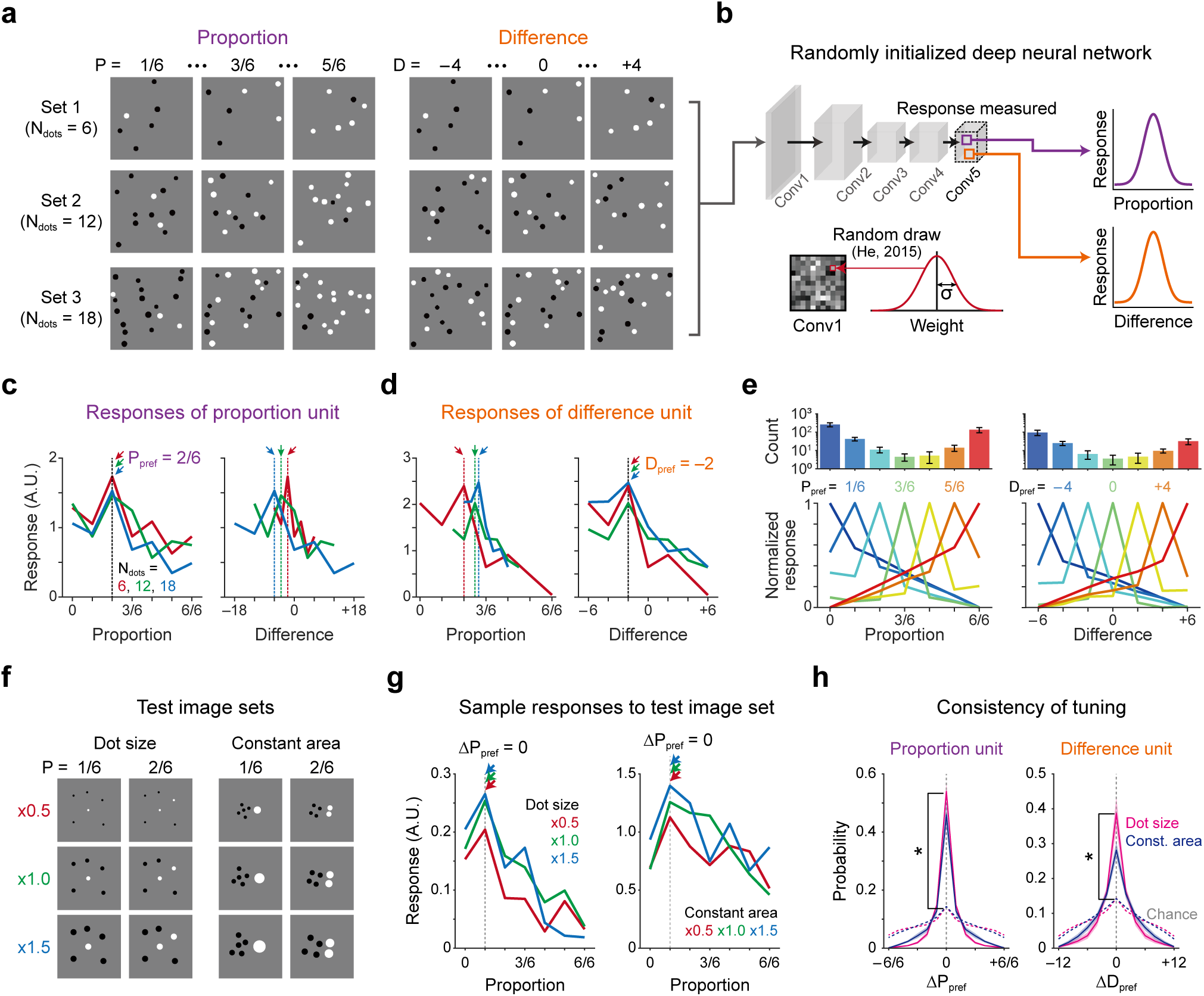
Units encoding quantity proportion and difference spontaneously arise in an untrained neural network. (a) Example of the stimuli used to find proportion- and difference-selective units in the network. Images were designed to represent seven different levels of proportion (difference) with black and white dots (200 images per condition), with three different total numbers of dots (N_dots_ = 6, 12 and 18). (b) Architecture of the untrained AlexNet, where the weights in each convolutional layer were randomly initialized^54^. Unit responses were calculated from the images shown in **a**. (c-d) Sample responses of proportion and difference units. Each line indicates an average response to the stimulus sets with different N_dots_. Each arrow indicates the peak position of the tuning curve. (c) A proportion unit response shows the same preferred proportion (P_pref_) on the proportion axis (left) but not on the difference axis (right). (d) A difference unit has the same preferred difference (D_pref_, right) but not on the proportion axis (left). (e) Number of proportion and difference units, and their average tuning response. Top: Distribution of preferred proportions and differences in Layer 5 (Mean ± S.D. from N_net_ = 20). Bottom: Population tuning curves of proportion and difference units. (f) Example of the stimuli used in the test for tuning consistency. Images were designed to have different dot sizes (left) or to have the constant area of white and black dots (right). (g) The sample response to the test image sets. P_pref_ was compared across the conditions. (h) Consistent tuning in each test set. The distributions of ΔP_pref_ and ΔD_pref_ were sharply concentrated to zero in all conditions, which was significantly higher than a chance distribution (*P < 1.26 × 10^-15^, paired t-test, N_repeat_ = 20 for control, N_net_ = 20). See Supplementary Figure S1.

### Spontaneous emergence of proportion- and difference-selective units in untrained networks

Remarkably, we found units that selectively respond to proportional differences and to different stimulus quantities in this randomly initialized network (Figs. 1c and d, N_unit_ = 474 ± 64 for proportion, N_unit_ = 179 ± 34 for difference in the Conv5 layer; see Materials and Methods for details). The “proportion-selective” units showed a maximum response at a particular stimulus proportion value regardless of the total number of dots (Fig. 1c, left, preferred proportion P_pref_ = 2/6 for N_dots_ = 6, 12, and 18). These proportion-selective units show different peak positions when their tuning curves are plotted on a “difference” scale on the x-axis (Fig. 1c, right), demonstrating that they are selective to the same proportion but not to the same differences in dot numbers. Similarly, we also found units selective to differences in dot numbers but not to proportion (Fig. 1d), confirming that there exist two distinct populations of proportion- and difference-selective units. We observed that the distributions of P_pref_ and D_pref_ covered the entire range of the presented images, and the population average tuning curves for those with the same preferred peak value are sharply tuned (Fig. 1e). We also observed that units preferring the smallest and the largest values of P_pref_ and D_pref_ were more frequently observed than those preferring mid-range values (Fig. 1e, top), similar to our previous observation of units with spontaneous visual number tuning^35^.

Next, to test whether proportion- and difference-selective units can maintain their tunings under variations of the visual parameters of the stimulus, we measured unit responses using controlled images in which the parameters of the dots were varied (Fig. 1f, Supplementary Figure S1). Particularly, considering the possibility that proportion and difference in a stimulus co-vary with the area of the dots, we investigated unit responses with image sets with 1) each dot variously sized (Fig. 1f, left) and 2) dot size inversely proportional to the total dot number so that the total stimulus area remained constant (Fig. 1f, right). We confirmed that the peak position of the tuning curve is maintained under these control simulations for both proportion and difference units (Fig. 1g, Supplementary Figure S1). To quantify the consistency of tuning, we measured changes in the preferred peak values of tuning curves for each stimulus set pair. The distributions of both ΔP_pref_ and ΔD_pref_ peak sharply at zero, and the peak amplitude was significantly higher than that of a chance level (Fig. 1h, *P < 1.26×10^-15^, paired t-test, N_repeat_ = 20 for control, N_net_ = 20; see also Supplementary Figure S1), demonstrating that the tuning of proportion and difference units is fairly robust. Similarly, we also confirmed that both units retain their tuning robustly while varying both the total number of dots and the geometric distribution of the dots (Supplementary Figure S1).

### Visual quantity comparison by proportion and difference units

With the notion that quantity comparisons in the brain may be executed by neural activities tuned to proportion and difference, we tested whether proportion or difference units enable the network to compare quantities represented by a pair of images, as observed in animals and human behavior^26,27,38^. Following previous approaches^35,39^, we measured population responses to a given pair of images and trained a support vector machine (SVM) using these activities to select an image representing the larger value of proportion or difference (Fig. 2a). Then, we tested whether the trained SVM can perform the comparison task with novel image pairs (Figs. 2a-b, see Supplementary Figure S2 for a matching task). We found that both the proportion and difference values of an image pair can be successfully compared by an SVM when trained with proportion and difference units, as distinguished from an SVM trained with non-selective units (Fig. 2b; *P = 1.13×10^-24^ for proportion comparison, *P = 3.07×10^-16^ for difference comparison, paired t-test, N_net_ = 20). Notably, the comparison performance increases as the distance between the pairs of proportions increases (Figs. 2c-d), reproducing the *distance effect* observed in humans and monkeys^26,27,38^. We also observed that the activity levels of selective proportion units were higher in correct trials than in incorrect trials, a finding also observed in proportion-selective neurons in a monkey experiment (Fig. 2e)^26^. These observations suggest that proportion and difference units emerged spontaneously in our untrained networks share common characteristics with selective neurons observed in animal brains.

**Figure 2.**
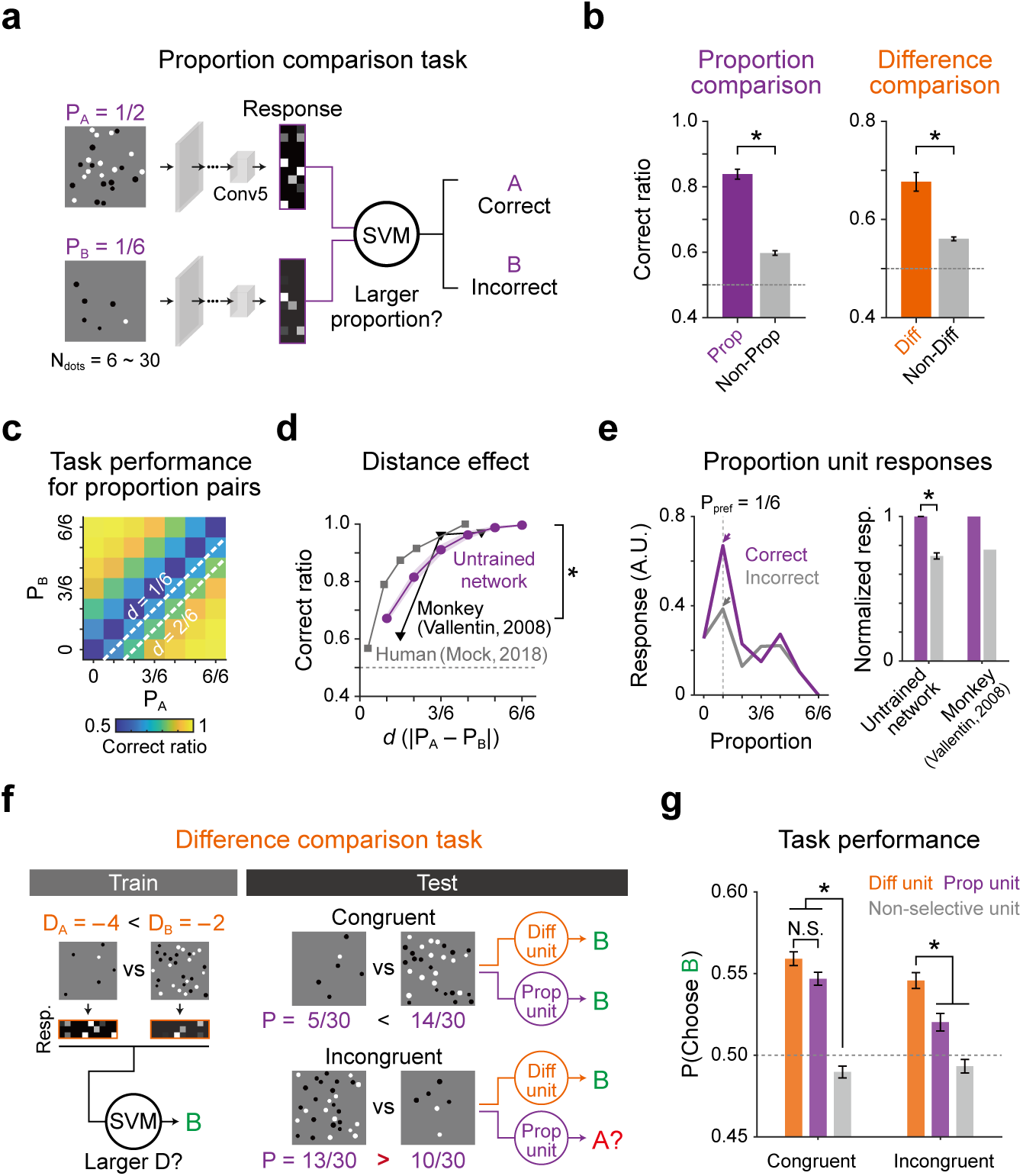
Proportion and difference units in an untrained network enable the network to compare visual quantities. (a) Proportion comparison task. The SVM was trained to choose an image with a larger proportion value between the two images. (b) Performance on the proportion- and difference-comparison tasks (*P = 1.13×10^-24^ for proportion comparison, *P = 3.07×10^-16^ for difference comparison, paired t-test, N_net_ = 20). (c-d) Performance across different pairs of proportions. The performance increases as the proportion distance (*d* = |P_A_-P_B_|) increases (distance effect, *P = 1.30×10^-28^, paired t-test, N_net_ = 20), similar to human and monkey data^26,38^. (e) Left: Sample response of a proportion unit during correct and incorrect trials. Each arrow indicates the response to a stimulus with the preferred proportion. Right: A greater response was observed during correct trials, similar to that of primates^26^ (*P = 9.90×10^-23^, one-sample t-test). (f) Difference comparison task using proportion and difference units. A pair of images was categorized as either a congruent and incongruent pair (see Materials and Methods for details). (g) Average task performance. Both units successfully compared congruent image pairs, but difference units outperform proportion units or non-selective units for incongruent pairs (*P < 2.23 × 10^-308^ for congruent pairs, *P < 5.27 × 10^-4^ for incongruent pairs; one-way ANOVA with Tukey’s *post-hoc* test).

While proportion-selective neurons were observed experimentally in previous animal studies^26,27^, difference-selective neurons have not yet been reported in experiments. Theoretically, it is possible to compare any arbitrary quantity only using one type of tuning, e.g., proportion selectivity, because proportion and difference values are positively correlated in most conditions. However, both types of tuning, i.e., proportion and difference selectivity, may be required under such conditions, particularly when visual quantities represent an incongruent relationship between proportion and difference values. To simulate the distinct contribution of proportion and difference units under this circumstance, we conducted an additional test in which SVMs were trained to choose an image with the larger value of the difference between D = −4 and D = −2 using either proportion or difference unit responses (Fig. 2f). For this test, two types of test images sets were prepared: congruent and incongruent pairs of proportion and difference values (Fig. 2f). Specifically, in the congruent pairs, image B has larger difference and larger proportion values than image A. On the other hand, in the incongruent pairs, image B has a larger difference value but a smaller proportion value compared to those of image A (see Materials and Methods for details). We found that both proportion units and difference units can successfully compare congruent pairs (Fig. 2g, left), whereas difference units outperformed proportion units when comparing incongruent pairs (Fig. 2g, right, *P < 5.27×10^-4^, one-way ANOVA with Tukey’s *post-hoc* test, N_repeat_ = 50, N_unit_ = 600). This is understandable because only proportion units cannot properly compare difference values under this controlled condition (Figs. 1c-d, see Supplementary Figure S3 for proportion comparison task). These results imply that both difference units and proportion units are required for consistent performance of quantity comparisons in the brain, suggesting that difference-selective neurons exist and can be found in biological brains.

### Selective responses from the feedforward summation of increasing and decreasing unit activities

To understand how proportion- and difference-selective units originate from a randomly initialized network without training, we investigated the detailed mechanism of development by initially counting the number of comparison units emerged in each layer (Fig. 3a)^36^. Among five convolutional layers, proportion and difference units were most commonly observed in deeper layers (Layers 3–5) but were rarely observed in early layers (Layers 1-2, see Supplementary Figure S4). Based on the observation that selective units start to emerge in Layer 3 (Fig. 3a, Layer 2 vs. Layer 3, *P < 9.99×10^-9^ (proportion), and *P < 9.99×10^-9^ (difference), one-way repeated measure ANOVA with Tukey’s *post-hoc* test), we assumed that the developmental mechanism of comparison units can be explained by the organization of connections between Layer 2 and Layer 3. Extending the notion from our previous study regarding the emergence of quantity selective units in untrained hierarchical networks^35^, we hypothesized that proportion and difference tunings also emerge from the feedforward summation of increasing and decreasing activities from early layers (Fig. 3b). Specifically, we introduced a summation model^35^ (Fig. 3b) which suggests that selective neural responses emerge from the weighted sum of monotonically increasing and decreasing activities from the previous layers. To validate this scenario, first we examined whether units of monotonically increasing or decreasing responses to variations of proportion values exist in our model network. Indeed, we found such units in our untrained networks; the activity of each unit monotonically increases or decreases as the corresponding proportion in the stimulus increases regardless of the total number of dots (Fig. 3c). Notably, these units were mostly observed in Layer 2 (36.5 ± 3.42% of Pool2 units for proportion, 13.0 ± 1.88% for difference) among the five layers, suggesting these units can be building blocks for proportion-selective units in the subsequent layers.

**Figure 3.**
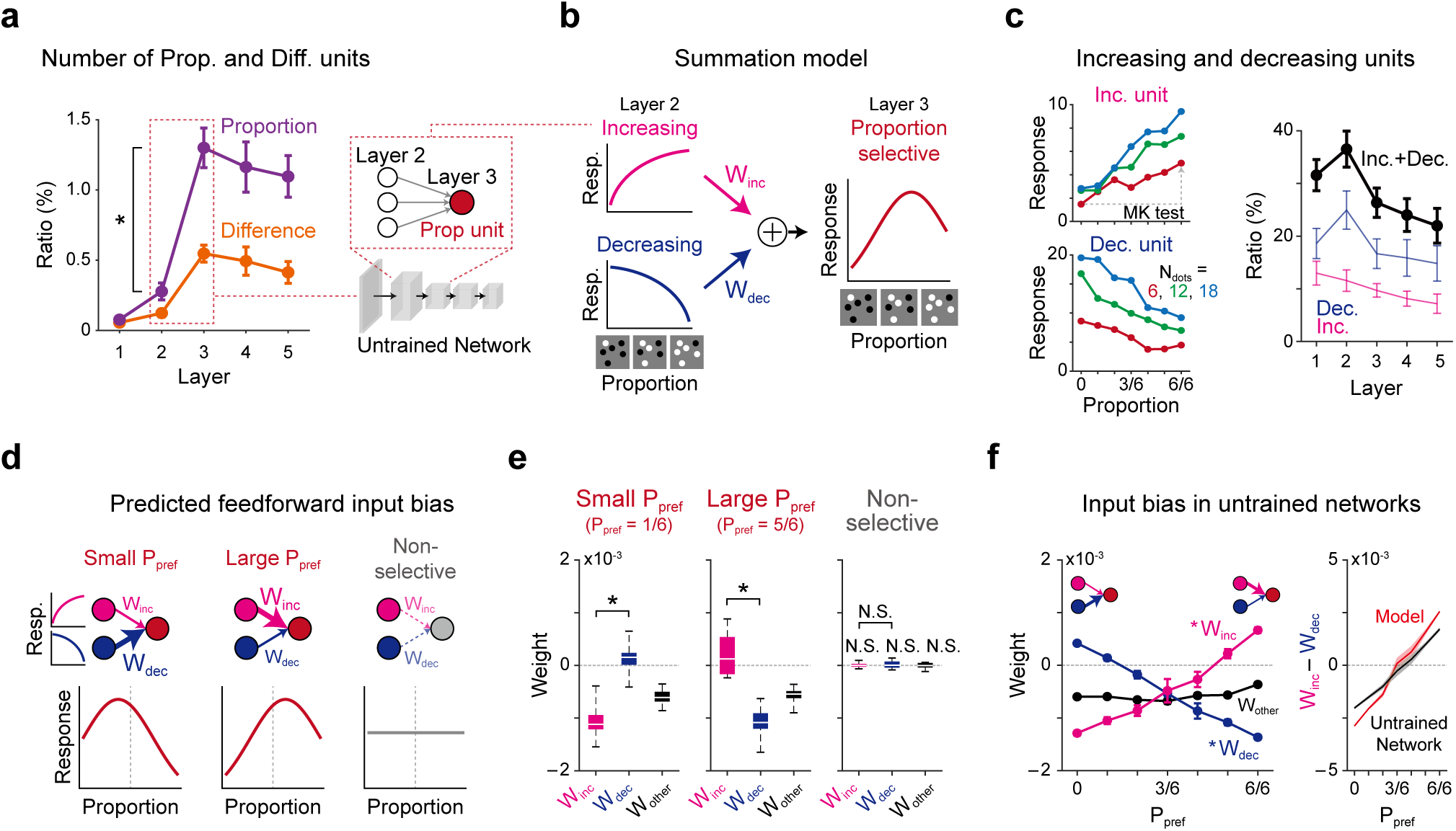
Theoretical model suggests that the summation of increasing and decreasing responses from the previous layer explains the spontaneous emergence of quantity-comparison units. (a) Numbers of two comparison units in each layer in untrained networks (N_net_ = 20). Note that both proportion and difference units start to emerge largely from Layer 3 (Layer 2 vs. Layer 3, *P < 9.99×10^-9^, one-way repeated measure ANOVA with Tukey’s *post-hoc* test). See Supplementary Figure S4 for the layer-wise tuning properties. (b) Summation coding model^35^. To generate a proportion-selective unit in Layer 3, the source units in Layer 2 in which the responses are increasing or decreasing should be summed. (c) Increasing and decreasing units. An increasing/decreasing unit shows a monotonically trending response along the proportion axis regardless of N_dots_ (left, P < 0.05, Mann-Kendall test). Numbers of increasing and decreasing units across layers are shown (right). (d) Predicted the input bias from the summation model. The model predicts that proportion unit with a low P_pref_ is receives stronger inputs from the decreasing unit (left), and vice versa for a unit with a high P_pref_ from an increasing unit (middle). The model also predicts that non-selective units have no bias (right). (e) Observed weight distribution from increasing and decreasing units to units in Layer 3 of untrained networks, consistent with the model prediction (left, *P = 7.37×10^-12^, middle, *P = 2.77×10^-11^, paired t-test, N_net_ = 20, N.S., P > 0.05). (f) Average W_inc_, W_dec_, and W_other_ of proportion units across different preferred proportions. Left: The increasing units (magenta, W_inc_) impart high weights to large P_pref_ units but low weights to small P_pref_, whereas decreasing units (blue, W_dec_) show the opposite trend (*P = 3.97×10^-4^, Mann-Kendall, N_net_ = 20). Right: Average weights estimated from the generation model (See Supplementary Figures S5-6).

Next, we examined whether core predictions of the summation model are verified in the current system. First, strong feedforward projections from increasing and decreasing units compose proportion units (Fig. 3b). Second, more specifically, proportion units with small P_pref_ values receive stronger inputs from decreasing units than from increasing units (Fig. 3d, left), while units with large P_pref_ values receive stronger inputs from increasing units (Fig. 3d, middle). Third, non-selective units receive unbiased input from both increasing and decreasing units (Fig. 3d, right). To validate this scenario, we analyzed the weight values between each unit in Layer 3 and the increasing and decreasing units in Layer 2 (W_inc_ and W_dec_, respectively). We found that the observed weight distributions accord closely with our model prediction — proportion units with small P_pref_ values receive significantly stronger inputs from decreasing units (W_dec_) than from increasing units (W_inc_) (Fig. 3e, left, W_inc_ < W_dec_, *P = 7.37×10^-12^, paired t-test, N_net_ = 20) and units with large P_pref_ values receive stronger inputs from increasing units (Fig. 3e, middle, W_inc_ > W_dec_, *P = 2.77×10^-11^, paired t-test). As a result, the feedforward connectivity weight for units with different P_pref_ values demonstrates a strong positive correlation between W_inc_ and P_pref_ and a negative correlation between W_dec_ and P_pref_, as expected from the model (Fig. 3f, *P = 3.97×10^-4^ for both W_inc_ and W_dec_, Mann-Kendall test, N_net_ = 20). In contrast, we found that non-selective units in Layer 3 receive unbiased weights from both increasing and decreasing units in Layer 2 (Fig. 3e, right). Given that all of these weights are randomly sampled from an identical Gaussian distribution from the random initialization, this result suggests that a small amount of bias caused by a random sampling can destine the tuning profile of selective units.

Notably, we found that the average weight of all feedforward projections to a proportion unit is negatively biased (Fig. 3f, one-sample t-test, P < 1.27×10^-10^ for all proportions), presenting strong tonic inhibition^40,41^, while such bias was not observed in the projections to non-selective units (Fig. 3e, right). To understand this unexpected finding, we introduced a simple model in which negative bias (tonic inhibition) in feedforward projection (mostly from W_other_) sharpens the tuning curve of a selective unit (Supplementary Figure S5). From this model simulation, we confirmed that the degree of proportion selectivity was correlated with the amount of negative bias in the weights. Further, we confirmed that a manually decreased W_other_ can sharpen the proportion-selective response in the model. We found a similar mechanism in the difference-selective units (Supplementary Figure S6).

Next, we tested whether comparison units can be generated from the random wiring of the observed monotonically increasing and decreasing units (Supplementary Figures S7a-c). We built a simple two-layer model in which the input layer is composed of the Layer 2 unit activities observed in untrained networks (Fig. 3c) and where the units in the output layer receive feedforward inputs from a randomly weighted sum of input layer activities. As expected, units selectively tuned to all possible proportions were generated (Supplementary Figure S7c) from these random wirings. In addition, we found that P_pref_ estimated from the observed tuning curve in the model was also significantly correlated with W_inc_ and W_dec_ (Fig. 3f), as previously observed in untrained networks. Furthermore, when the wiring strength from the increasing units (W_inc_) was manually increased, the number of proportion units with a large P_pref_ value increased, whereas when the wiring strength from the decreasing units (W_dec_) was increased, the number of proportion units with a small P_pref_ value increased (Supplementary Figure S7d-g). These model simulations demonstrate that proportion- and difference-selective units can emerge from the random wiring of increasing and decreasing unit activities that arise spontaneously in untrained networks.

Lastly, to verify the key assumption of the model that increasing and decreasing unit activities in Layer 2 are building blocks for the emergence of comparison units, we examined the changes of the network when units in Layer 2 are ablated (Supplementary Figures S7h-j). We compared the number of proportion (difference) units observed under three conditions: 1) a normal condition with no ablation, 2) with all increasing and decreasing units in Layer 2 ablated, and 3) with all Layer 2 units except increasing and decreasing units ablated. As predicted by the model, the ablation of increasing/decreasing units significantly modulates the tuning curve of the comparison units so that they lose their selectivity. As a result, the number of comparison units was significantly reduced (Supplementary Figures S7i-j). In contrast, the number of comparison units was increased when only increasing/decreasing units were connected and all other units were ablated. These results provide further support of the model and show that increasing and decreasing units are the source of the proportion/difference units that emerge spontaneously.

### Nonlinearity pattern of source units and corresponding type of selectivity in target units

Given that both types of comparison units emerge from a universal mechanism, a random summation of the increasing/decreasing units from an identical unit population, we examined how the two distinct types of measures, proportion and difference of quantities, can develop. A possible scenario from a mathematical model is that slightly different profiles of the nonlinearity of source unit activity that look similar but are described by different mathematical functions of quantity scaling can generate distinct measures of proportion or difference in target units (Fig. 4a). Specifically, for different sets of the total number of dots of the stimulus, a tuning curve composed of increasing and decreasing “power” functions has the same profile on the x-axis scaled as the “proportion” value of quantity (Fig. 4a, top). On the other hand, a tuning curve composed of increasing and decreasing “exponential” functions has the same profile on the x-axis scale as the “difference” value of quantity (Fig. 4a, bottom; see Supplementary Notes for a mathematical proof). This simple mathematical model provides a theoretical framework by which to understand the observed results of distinct tunings from a universal mechanism of development.

**Figure 4.**
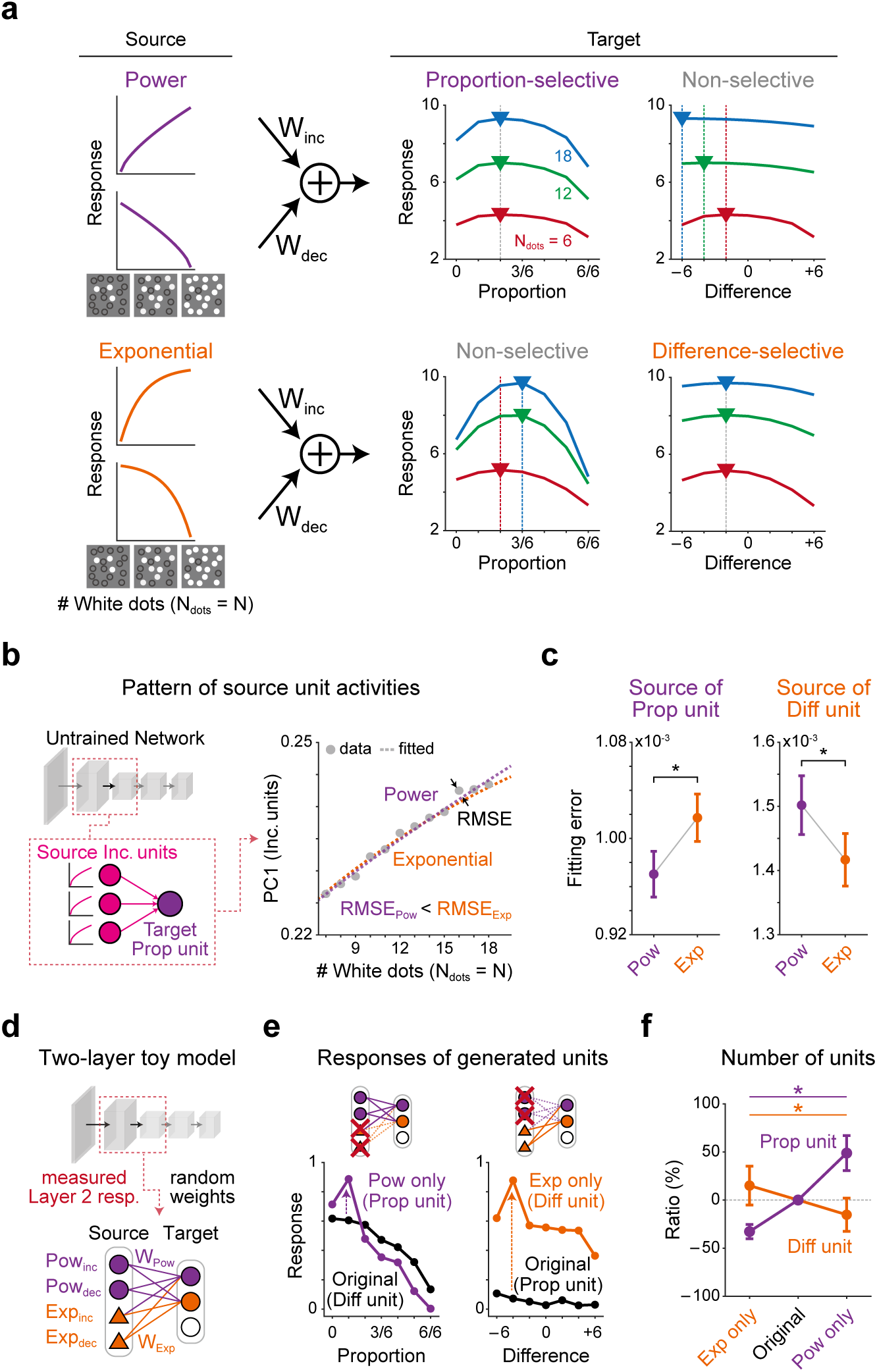
Summation of the power-like function generates proportion units, and summation of the exponential-like function generates difference units. (a) Mathematical model of the proportion and difference unit-generation mechanism. Top: The summation of power-increasing and -decreasing units have an identical peak in all proportion sets but misaligned peaks across difference stimulus sets. Bottom: The summation of exponential-increasing and -decreasing units have misaligned peak positions along the proportion axis but the same peaks on the difference axis. (b) Functional form of increasing source units. As suggested in **a**, the source units’ increasing responses were plotted against the number of white dots and the principal component of the increasing responses was calculated. The first PC component was fitted to either a power function or an exponential function and the root-mean-squared error (RMSE) was calculated. (c) Fitting errors. Source units connected to proportion units were better fitted to a power function but source units connected to difference units were better fitted to an exponential function (*P = 5.59 ×10^-4^ and 4.86 ×10^-7^ for proportion and difference units; paired t-test, N_net_ = 20). (d) Toy model for proportion and difference units. (e) Sample generated units. A proportion unit was generated when all exponential units were ablated (left) and a difference unit was generated when all power units were ablated (right). (f) The number of proportion and difference units in each unit ablation condition. The number of proportion units that emerges decreases whereas difference units increase when the power-response units are ablated, and vice versa when exponential-response units are ablated (*P = 3.36×10^-23^ for proportion units, 3.65×10^-7^ for difference units; one-way repeated measure ANOVA with Tukey’s *post hoc* test).

To validate this scenario, we examined the connectivity between the proportion and difference units in Layer 3 and the increasing/decreasing source unit activities in Layer 2 (Figs. 4b-c). First, we classified all increasing/decreasing source activities as “power-like” or “exponential-like” using their first principal component (PC1, Fig. 4b, left), which explains 98% of the variance of the increasing/decreasing responses (see Materials and Methods for details). We then fitted each pattern of PC1 to either power or exponential functions (Fig. 4b, right). We found that PC1 of the proportion-source units was much better fitted to a power function than to an exponential function (Fig. 4c, left, RMSE_Pow_ < RMSE_Exp_, *P = 5.59×10^-4^, paired t-test, N_net_ = 20, RMSE represents the root-mean-square error). In contrast, difference-source units were better fitted to an exponential function, agreeing with the predictions of the model (Fig. 4c, right, RMSE_Pow_ > RMSE_Exp_, *P = 4.86×10^-7^, paired t-test, N_net_ = 20; see Materials and Methods for details).

Next, we examined changes in the comparison units when these power-like or exponential-like source units are ablated (Figs. 4d-f, see Supplementary Figure S8). We built a two-layer toy model in which the input layer is composed of power-like or exponential-like activities in Layer 2 and where the units in the output layer receive feedforward inputs from the randomly weighted sum of the input layer activities (Fig. 4d). Then, we undertook the ablation of each source unit group and examined if this would affect the target unit activities (Fig. 4e). Remarkably, when all exponential-like source units were ablated so that only power-like source units remained connected, the proportion-selective unit activities were strengthened while the difference-selective activities were suppressed, as predicted. Moreover, some difference-selective units even transformed and became proportion-selective units in this case (Fig. 4e, left). In contrast, when all power-like source units were ablated, difference-selective unit activities were strengthened while proportion-selective activities were suppressed (Fig. 4e, right; see also Supplementary Figures S8f-g). Overall, the ratio of proportion units increased when only power-like inputs were connected and decreased when only exponential-like inputs were connected (Fig. 4f, purple, *P = 3.36×10^-23^), significantly different from that of the original condition. As predicted, the opposite trend was observed for difference units — “exponential-like only” inputs (power-like units ablated) generated more difference-selective units and “power-like only” inputs (exponential-like units ablated) suppressed difference-selective responses (Fig. 4f, orange, *P = 3.65×10^-7^ for difference units, one-way repeated measure ANOVA with Tukey’s *post hoc* test, N_net_ = 20). These results suggest that the two distinct types of measures, proportion and difference selectivity, develop from slightly different profiles of the nonlinearity of the source activity.

## Discussion

In this study, we showed that the capability of comparing visual quantities emerges spontaneously in untrained deep neural networks in the complete absence of learning. We found two distinct tunings, proportion- and difference-selectivity, in a randomly initialized AlexNet, which are invariant to low-level image features. These selective neural responses enable the network to perform quantity comparison tasks, reproducing the behavioral and neuronal characteristics observed in animal experiments. Our theoretical model demonstrates that these quantity comparison functions can originate from a combination of monotonically increasing and decreasing neural activities that arise spontaneously from random feedforward wiring in a hierarchical network model. Lastly, the model suggests that two distinct types of comparison functions, proportion- and difference-selectivity, can be separated by a slight difference in the nonlinearity of the preceding response profile.

One crucial finding in the current study is that two distinct types of comparison functions, proportion- and difference-selectivity, are developed spontaneously in untrained networks (Figs. 1c-e). A series of validation processes here proved that the observed tunings are selectivity type for proportion and difference senses that are invariant to low-level image features (Figs. 1f-h), with both types emerging very robustly in randomly initialized neural networks. Notably, the proportion sense observed in untrained networks shares the previously reported core characteristics of neural tunings in primate brains^26^. The proportion-selective units in our randomly initialized AlexNet enabled the network to compare two given proportion values (Figs. 2a-b) with a comparison accuracy well described by the distance effect (Fig. 2d), similar to earlier in primates^26,38^. Greater responses of proportion-selective neurons for the correct choice in primate experiments were also replicated by the units in untrained networks (Fig. 2e). Although the proportion units observed here may not be an exact homologue of proportion-selective neurons in the prefrontal cortex of primates, our results suggest a probable scenario for understanding the early development of quantity comparison sense in the brain.

On the other hand, neurons that selectively respond to quantity differences have not yet been reported in biological brains, although it is widely observed that animals as well as humans can utilize difference computations, or subtraction, between two visual quantities when making decisions^42,43^. In our observation in untrained neural networks, difference-selective units are predicted to be found along with proportion-selective units, probably in similar anatomical hierarchies (Fig. 3a) and with a common mechanism of development (Fig. 3). Specifically, our model simulation results suggest that difference-selective neurons are more difficult to find than the proportion-selective neurons, for a couple of reasons; 1) the probability of spontaneous emergence is lower than that of proportion units (Fig. 3a), and 2) in single-neuron experiments, they are not readily distinguishable from proportion-selective neurons unless careful validations take place using multiple sets of quantities representing distinct values of differences and proportions (Figs. 1a-d). Nevertheless, our results suggest that difference-selective neurons can be observed experimentally, which may provide a fundamental basis of the innate arithmetic function in the brain, along with the proportion-selective neurons previously reported.

On the mechanism underlying the spontaneous emergence of quantity-comparison units, we showed that the summation model proposed in our previous study to explain the emergence of number sense in untrained networks^35^ can also explain the emergence and segregation of two types of tunings, proportion and difference selectivity (Figs. 3a-b). A new crucial finding is that a slight difference in the nonlinearity of the source unit characteristics can lead to the emergence of completely different types of tunings, such as difference and proportion selectivity (Fig. 4). Considering that the feedforward wiring in young brains have a variety of connection profiles and weight distributions^44^, similar to the that of randomly initialized deep neural networks, the current result suggests that a wide repertoire of neuronal functions can originate from a universal development mechanism but then diverge due to certain determinant factors, such as the degree of nonlinearity or the response threshold. This is a plausible scenario with regard to the origin of diverse innate functions in randomly wired young brain circuits. Our series of analyses (Figs. 3d-f, Supplementary Figures S5-7) provide supporting evidence that a biased sampling of feedforward projections in which the weight of each wiring is drawn from an identical Gaussian distribution is the core component to generate distinct neuronal tunings without learning. Another interesting observation in our model simulation, in addition to the biased weight projection from increasing and decreasing units, is the constant negative bias of the total weights from the entire set of feedforward source units (Figs. 3e-f). Our further analysis suggests that this consistent negative bias can further sharpen the tuning (Supplementary Figure S5), similar to the selectivity modulation mechanism in sensory neurons, often driven by inhibitory interneurons^40,41^. Overall, the current observations from the artificial neural networks may provide insight into how complicated neural mechanisms in the brain can be studied and understood using computational approaches with a simplified model neural network.

Definite advantages of a computational model simulation to study cognitive functions in neural networks are that controlled datasets can be tested infinitely and that any circuit components are readily accessible and testable either on a network scale or at a single-neuron level, which is very challenging in biological brain studies. Taking advantage of a computational model, we examined a massive number of neural responses (650,080 units in total of Layers 1-5) to a huge amount of image stimulus sets (8,400 images in total for proportion and difference sets) with variations of the control features. Moreover, we attempted to manipulate relevant circuit structures to test our key hypothesis — dissect the weights between units in an untrained network (Figs. 3d-f, Supplementary Figures S5-6), artificially generate a tuned unit using a theoretical model (Supplementary Figures S7a-g), and ablate a group of units to reveal the necessary conditions for tuned responses (Supplementary Figures S7h-j). With a series of these model validations, we successfully demonstrated that the biased summation of feedforward afferents is sufficient for the spontaneous emergence of quantity-comparison units in an untrained network. Although AlexNet may not be a completely plausible model of the ventral visual pathway, our result provides a possible explanation of how a primitive form of a quantity comparison system arises from random feedforward connections in untrained neural networks. While visual experience and learning can significantly contribute to the development of quantity perceptions in adult brains^20,45,46^, our result implies that neuronal functions for quantity comparisons emerge prior to learning in the young brains of infants or newborn animals. Along with our recent studies of the emergence of cognitive functions in untrained neural networks^35,36^, our findings here provide insight into how cognitive functions arise initially in both biological and artificial neural networks^47–53^.

## Methods

### Neural network model

AlexNet^34^ was used as a representative model of a convolutional neural network. This network consists of two parts: feature extraction and classification networks. In detail, the feature extraction network consists of five convolutional layers with rectified linear unit (ReLU) activation and a pooling layer, while the classification network has three fully connected layers. The detailed parameters of the architecture were sourced from Krizhevsky et al. (2012), which provided the models for V4 and IT.

To investigate how proportion- and difference-selective units can arise spontaneously, randomly initialized networks were examined. For the untrained AlexNet, the weights of each convolutional layer were initialized from a Gaussian distribution with a zero mean and the standard deviation set to balance the strength of the input signals across convolutional layers (bias = 0)^54^. All simulations were conducted for 20 networks.

### Stimulus dataset

The stimulus sets (Fig. 1a) were designed based on earlier work^35,39^. Briefly, images (size, 227 × 227 pixels) that contain several white and black dots were provided as inputs to the network. To ensure tuning for proportion (difference) under different total number of dots, we designed three stimulus sets (N_dots_ = 6, 12, 18) in which the numbers of white and black dots represent a proportion or a difference. For all sets, dots were located at random locations but with a nearly consistent radius (generated by a normal distribution; mean = 7, SD = 0.7 pixels). To compare the proportion units and difference units under the same statistical power, we used seven values each for proportion and for difference. The proportion was set to 0/6, 1/6, 2/6, …, 1 (step size: 1/6) and the difference was set to −6, −4, …, 6 (step size: 2). The number of images for each condition was set to 200.

Stimulus sets to test consistent tuning (Fig. 1f and Supplementary Figure S1) were designed similarly, but with the following constraints. First, the total number of dots varied (N_dots_ = 6, 12, 18, 24, and 30 dots). Second, the total area of the dots remained constant (500, 1000, and 1500 pixels^2^ for white and black dots, respectively) across different proportions (differences). Third, the size of each dot was varied (0.5 or 1.5 times). Finally, the convex hull of the total number of dots was fixed as a regular pentagon and the shape of each dot was determined to be that of a circle, a rectangle, an ellipse, or a triangle with an equal probability of each.

### Analysis of the responses of the network units

The responses of network units in the convolutional layer (after ReLU activation of the convolutional layer) were mainly analyzed. Similar to the method used to detect number-selective units in previous work^35^, a two-way analysis of variance (ANOVA) with two factors (proportion and stimulus set, difference and stimulus set) was used to define proportion- and difference-selective units. To detect proportion-selective units generating a significant change of the response across proportion but with an invariant response across stimulus sets (different total number of dots), a network unit was considered to be a proportion-selective unit if it exhibited a significant change for proportion (P < 0.05) but no significant change for the stimulus set or interaction between two factors (P > 0.05). To ensure the invariance of proportion tuning, selective units showing 1) the same peak positions across stimulus sets and 2) a high selectivity index (SI ≥ 0.4) were selected as proportion units, while the selectivity index was defined using the following equation:

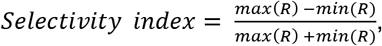

where *R* represents the average responses to proportion stimulus sets. In contrast, a network unit was considered to be non-selective (non-proportion unit) if it exhibited no significance for all factors. The same procedure was applied to detect difference-selective units.

The preferred proportion (P_pref_) was defined as the proportion that induced the largest response for each stimulus set. This also applies to the preferred difference (D_pref_). To determine the average tuning curves of all proportion units, the tuning curve of each unit was normalized by mapping the maximized response to 1 and was then averaged across units using the P_pref_ value as a reference point (Fig. 1e). The tuning curves were then normalized by mapping the minimized and maximized responses to 0 and 1, respectively. This was also done in the difference unit.

### Proportion (Difference) matching task for the network

The proportion/difference matching task was designed to test whether the selective unit activities observed in untrained networks would enable the network to classify the proportion or difference value of a given image (Supplementary Figure S2). For the proportion matching task, a sample stimulus (P = 0, 1/6, …, 1) was presented to the network and the resulting responses of the proportion units (Layer 5, Relu5) were recorded. Then, an SVM was trained with the responses of 120 randomly chosen proportion units (ten trials of random sampling for each untrained network) to classify the proportion value of the given image. To ensure invariance to the total number of dots, stimulus sets with the different total number of dots were used for training and test sets, respectively (e.g., N_dots_ = 6 for the training set, N_dots_ = 18 for the test set). As a control, a unit that does not exhibit a selective response to the proportion set (non-prop unit) was used for SVM training. The same procedure was applied to the difference comparison task (i.e., D = −6, −4, …, +6).

### Proportion (Difference) comparison task for the network

First, a vanilla version of the proportion/difference comparison task was designed to examine whether proportion (difference) units can sufficiently perform a task that requires an estimation of proportion from images (Fig. 2). For the proportion comparison task, a sample and a test stimulus (P = 0, 1/6, …, 1) were presented to the network and the resulting responses of the proportion units (Layer 5, Relu5) were recorded. Then, an SVM was trained with the responses of 120 randomly chosen proportion units (ten trials of sampling for each untrained network) to predict whether the proportion of the sample stimulus is greater than that of the test stimulus. To ensure invariance to the total number of dots, the total number of dots in each image was varied from 6 to 30. As a control, a unit that does not exhibit a selective response to proportion set (non-proportion unit) was used for SVM training. The same procedure was applied to the difference comparison task (i.e., D = −6, −4, …, +6).

Next, to reveal the distinct role of the two types of comparison units, comparison conditions were divided into two conditions: congruent and incongruent cases for difference and proportion comparison tasks (Figs. 2f-g, Supplementary Figure S3). In the difference comparison task, an image representing a larger difference can be compared with either an image representing a smaller difference and a smaller proportion value (congruent) or an image with a larger proportion value (incongruent). Image pairs were generated from four different conditions ((N_white_, N_black_) = (1, 5), (2, 4), (13, 17), or (14, 16), representing D = −4 or −2), resulting in three congruent pairs and an incongruent pair. The SVM was trained with both congruent and incongruent sets. The proportion comparison task was designed similarly (Supplementary Figure S3).

### Weight analysis between two convolution layers

The increasing (or decreasing) units were determined based on the Mann-Kendall test. To be a proportion-increasing (decreasing) unit, a unit needs to show significantly (P < 0.05) increasing (decreasing) responses by proportion across all stimulus sets. This also applies to difference-increasing/decreasing units.

To investigate the connectivity pattern between Pool2 increasing/decreasing units and Relu3 proportion (difference) units, Pool2 monotonic units within a convolutional filter of a specific proportion unit are considered to be connected with the corresponding weights (Supplementary Figure S7). Then, the weights between Pool2 monotonic units and proportion (difference) units with a specific preferred proportion (difference) were concatenated and their average was calculated in a single untrained network (Figs. 3e-f).

### Generation model for the weighted summation of increasing and decreasing units

A simple two-layer network simulation was designed to show that the summation of increasing and decreasing unit responses can reproduce the tuning responses of proportion or difference units. The tuning curve of a model output neuron (*R*) was defined by

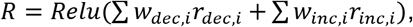

where *w_dec,i_* and *w_inc,i_* are the weight of the *i* th decreasing or increasing unit, respectively, and *r_dec,i_* and *r_inc,i_* indicate the corresponding tuning curves. The decreasing and increasing unit responses (*r*) were from the observed activities of the Pool2 layer of the untrained AlexNet. Estimated from the connection of the AlexNet, 526 decreasing units, 240 increasing units, and 1538 non-increasing/decreasing units were randomly sampled for the model simulation. The feedforward weight (*w*) was also randomly sampled from the Gaussian distribution estimated from the untrained AlexNet. In a trial, 64,896 output units (identical to the number of units in the Relu3 layer) were generated and 20 trials were performed for the simulation. The criteria used to detect proportion and difference units were identical to those used with the untrained networks.

### Mathematical model for proportion and difference units

To be a proportion (difference) unit, the peak positions of tuning curves across three stimulus sets (N_dots_ = 6, 12, 18) need to be aligned. To find a condition in which either a proportion or difference unit arises, we mathematically and numerically simulate the weighted sum when hypothetical increasing and decreasing the responses. In Figure 4, the power function and the exponential function were used for generating proportion units and difference units, respectively (see Supplementary Figure S8 for the detailed procedure).

### Connectivity analysis between power/exponential-like units and selective units

To determine the profile of increasing/decreasing responses connected to proportion/difference units, a principal component analysis (PCA) was applied to the monotonic responses of units connected to proportion or difference units, respectively. To determine whether PC1 shows power-like or exponential-like responses, the root-mean-square error (RMSE) between the fitted curve and the data curve in each case was compared. Because the response for stimulus set 1 (N_dots_ = 6) is identical, the RMSEs except for set 1 were compared (Fig. 4).

### Generation model for proportion and difference units

All procedures are identical to the generation model in Figure 3 but power-like and exponential-like responses were used as inputs. To classify the increasing (decreasing) units into two groups based on whether the unit response is the power-like or exponential-like type, we calculated the error between the normalized responses of each stimulus sets on the proportion axis and difference axis, respectively (Supplementary Figure S8). A unit showing a smaller error on the proportion axis than on the difference axis was determined as a power-like unit, and vice versa for exponential-like units.

### Statistical analysis

All statistical variables, including the sample sizes, exact P values, and statistical methods, are indicated in the corresponding text or figure legends.

## Code availability

MATLAB (MathWorks Inc.) with deep learning toolbox was used to perform the simulation and the analysis. The MATLAB codes used in this work are available at https://github.com/vsnnlab/Comparison.

## Acknowledgments

This work was supported by a grant from the National Research Foundation of Korea (NRF) funded by the Korean government (MSIT) (No. NRF-2022R1A2C3008991, NRF-2021M3E5D2A01019544, NRF-2019M3E5D2A01058328 to S.P., and NRF-2021R1C1C2007086 to W.C.) and by the Singularity Professor Research Project of KAIST (to S.P.).

## Author contributions

S.P. conceived of the project. H.L., W.C., and S.P. designed the model. H.L. and W.C. performed the simulations. H.L. and W.C. analyzed the data. H.L., W.C. and D.L. drafted the manuscript. H.L. and W.C. designed the figures.

S.P. wrote the final version of the manuscript. All authors discussed and commented on the manuscript.

## Competing interest declaration

The authors declare that they have no competing interests.

## Supplementary Information

**Figure S1.**
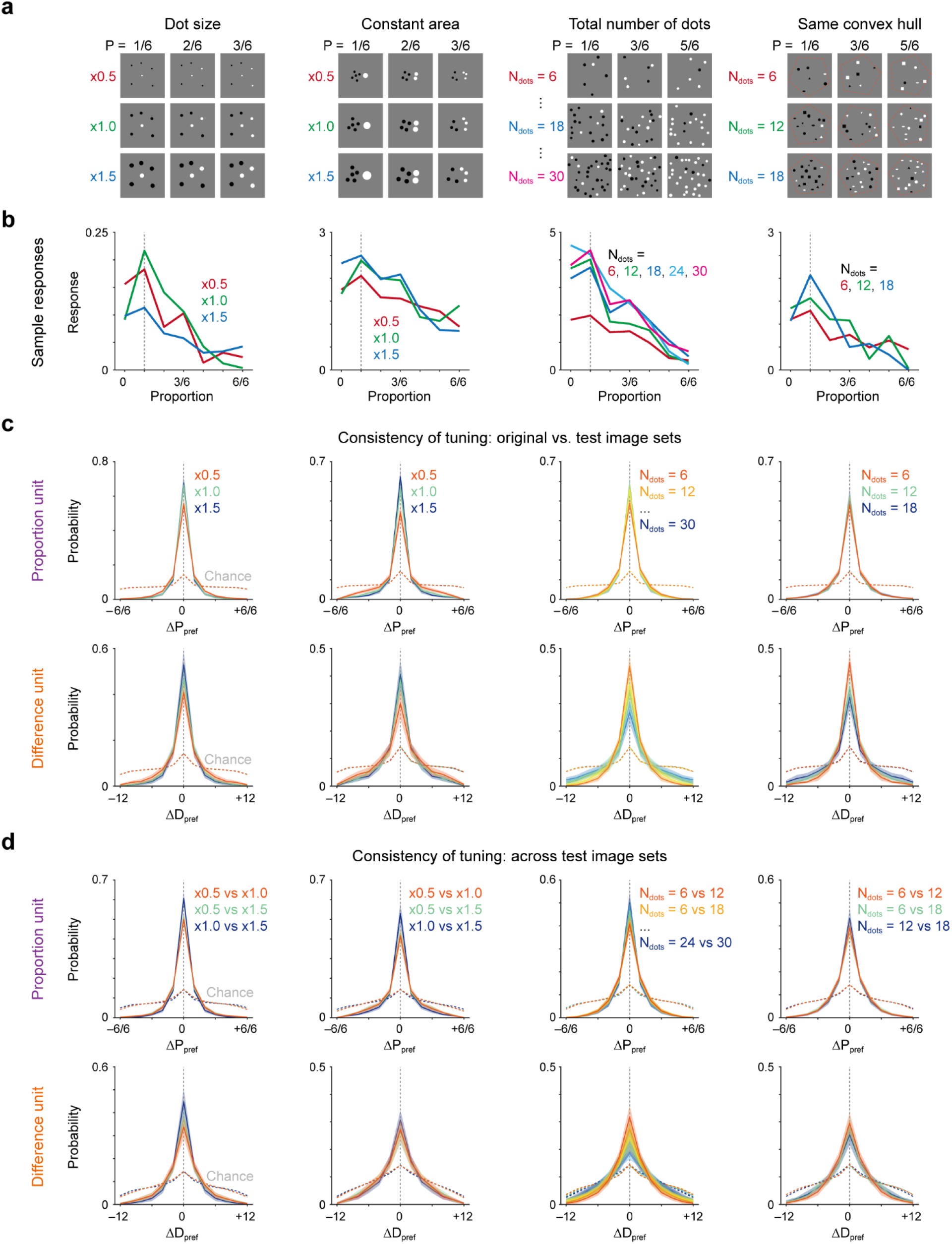
Consistent tuning of proportion and difference units. (a) Design of test image sets. Each set of images was designed to have 1) various dot sizes (Fig. 1f, left), 2) a constant total area of the dots (Fig. 1f, right), 3) various total number of dots (maximum of 30 dots), and 4) the same geometric distribution of the dots. (b) Sample unit responses to the test image sets in **a**. (c) Comparison of preferred proportions estimated from the responses to the original stimulus set and the test stimulus sets. The difference in P_pref_ (ΔP_pref_) between each pair of image sets was measured. In all conditions, ΔP_pref_ peaks sharply at zero and is significantly higher than the chance level. (d) Comparison of preferred proportions estimated from the responses to the test stimulus sets. Each panel uses a format identical to that in **c**.

**Figure S2.**
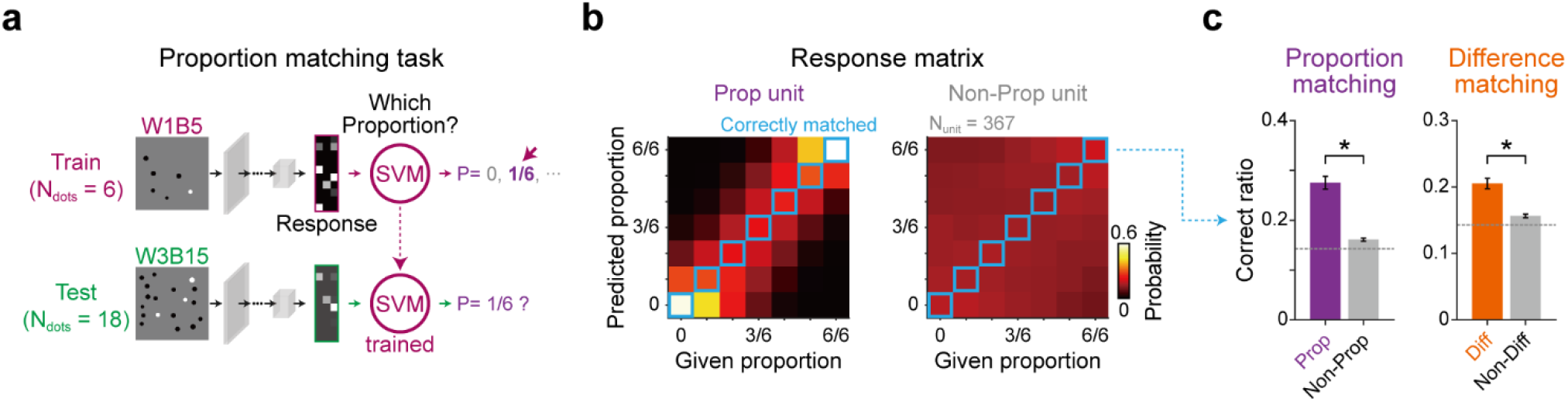
Proportion and difference matching tasks. (a) Proportion matching task. The SVM was trained to match the proportion value of a given image using the proportion unit responses. The population unit responses for images with different values of proportion with a fixed total number of dots (e.g., N_dots_ = 6) were measured and a support vector machine (SVM) was trained with these unit responses to select the correct value representing the given image. Then, the trained SVM was tested to perform the task with novel images in which the total number of dots differs from that used for training (e.g., N_dots_ = 18). (b) Response matrix of the SVM. The performance of the SVM was defined as the probability of correctly predicting a given proportion value (diagonals of the matrix). (c) Performance outcomes for the proportion- and difference-matching tasks. Note that both proportion and difference values can be correctly matched by unit responses, as the performance outcome of the SVM trained with proportion and difference units were significantly higher than those of the SVM trained with non-selective units (*P = 1.57×10^-19^ for proportion matching, *P = 5.72×10^-17^ for difference matching, paired t-test, N_net_ = 20).

**Figure S3.**
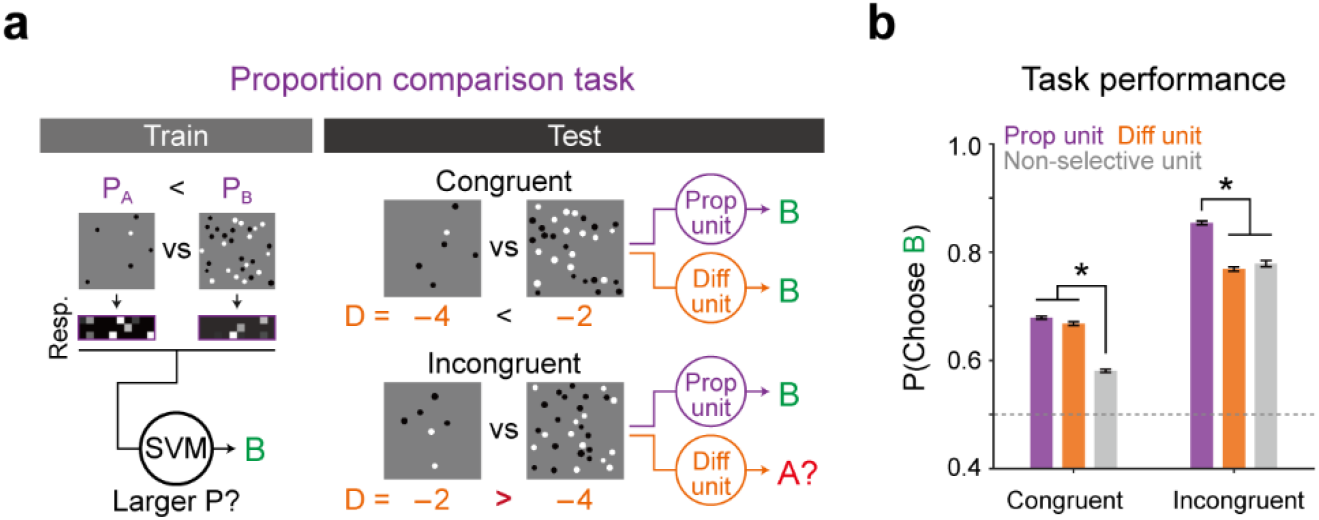
Proportion comparison task using proportion and difference units. (a) Design of the proportion comparison task. A pair of images is categorized into two classes: congruent and incongruent. In a congruent pair, an image with a larger proportion has a larger difference value, whereas an incongruent pair has a smaller difference. An SVM is trained using proportion unit responses and difference unit responses and is tested to compare the proportion of each pair of images. (b) Average performance of each type of unit. (left) When a congruent pair of images is given, both proportion and difference units can successfully compare the proportion values significantly better than non-selective units can (left, *P < 2.23×10^-308^ for congruent pairs, one-way ANOVA with Tukey’s *post-hoc* test). (right) When an incongruent pair of images is given, proportion units outperform difference units or non-selective units (right, *P < 2.23×10^-308^, one-way ANOVA with Tukey’s *post-hoc* test).

**Figure S4.**
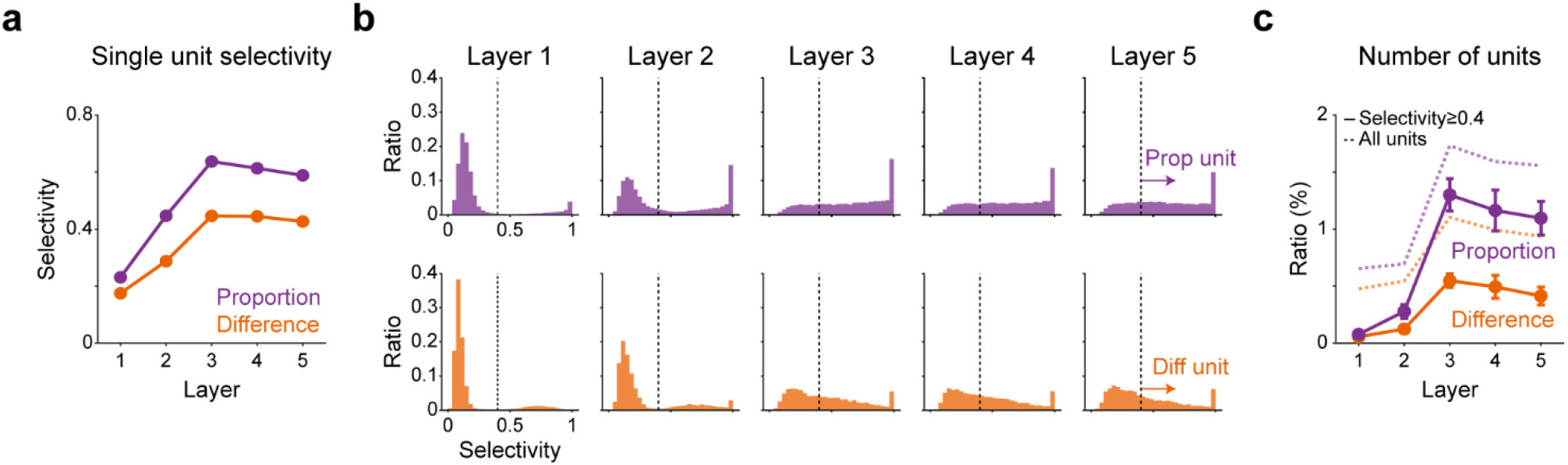
Layer-wise characteristics of proportion and difference units. (a) Single-unit selectivity of proportion and difference units across layers (see Materials and Methods for the definition of a selectivity index). The selectivity indices of proportion and difference units in the deep layers (Layers 3-5) are significantly higher than those in the shallow layers (Layers 1-2, P < 1.29×10^-8^ for a proportion unit, P < 1.02×10^-8^ for a difference unit, one-way repeated measure ANOVA with Tukey’s *post-hoc* test, N_net_ = 20). (b) Distribution of the single-unit selectivity of proportion and difference units across layers. In the shallow layers (Layers 1-2), the majority of units have low selectivity (less than 0.4) as opposed to that observed in deep layers (Layers 3-5). The black dotted line represents the threshold for determining selective units (Threshold value = 0.4). (c) The number of selective units across layers. Dotted lines represent the number of selective units without considering the threshold. Solid lines represent the number of selective units after thresholding.

**Figure S5.**
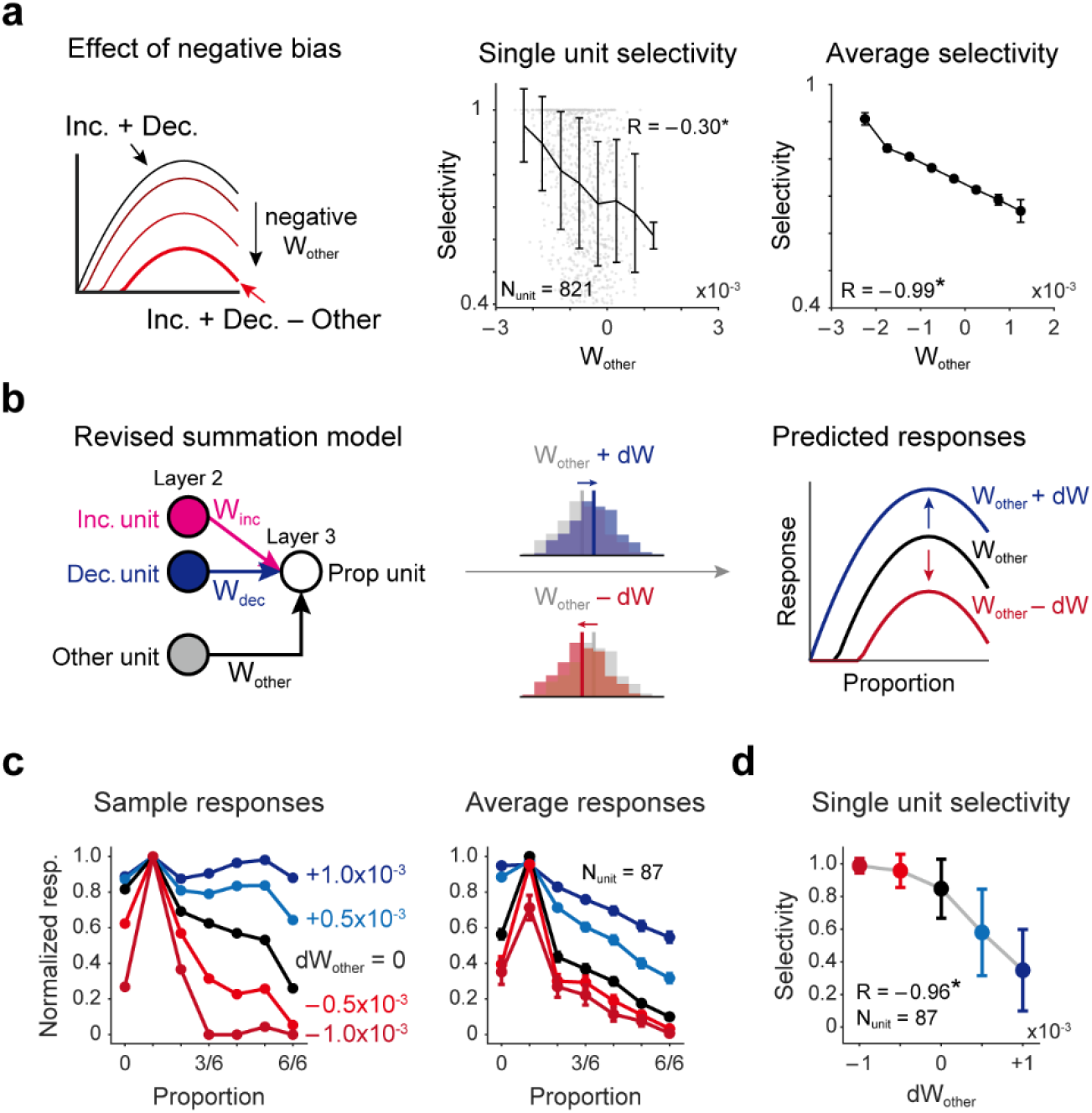
Effect of W_other_ on proportion tuning selectivity. (a) Effect of negative bias from non-increasing/decreasing units. The tuned response is initially generated by increasing and decreasing responses and is sharpened by the negative bias from other units (left). The observed proportion-selectivity is strongly correlated negatively with the average weights from other units (*P = 9.47×10^-19^, R = −0.30, between W_other_ and the single-unit selectivity, *P = 7.74×10^-6^, R = −0.99, between the W_other_ and average selectivity, Pearson correlation). (b) Revised summation model. The model hypothesizes that the weighted summation of increasing and decreasing units generates proportion (difference) tuning, with the bias from other units controlling the degree of tuning by modulating the response gain. Accordingly, higher selectivity is predicted when the value of W_other_ decreases. (c) Changes in the sample and the average unit response by the manipulation of W_other_. As the bias decreases and becomes negative (dW_other_ < 0, red curves), the tuning curves become sharpened, but they become flatter when positive biases are given (dW_other_ > 0, blue curves). (d) A negative correlation between the change of W_other_ and the single-unit selectivity (*P = 0.01, R = −0.96, Pearson correlation) is observed.

**Figure S6.**
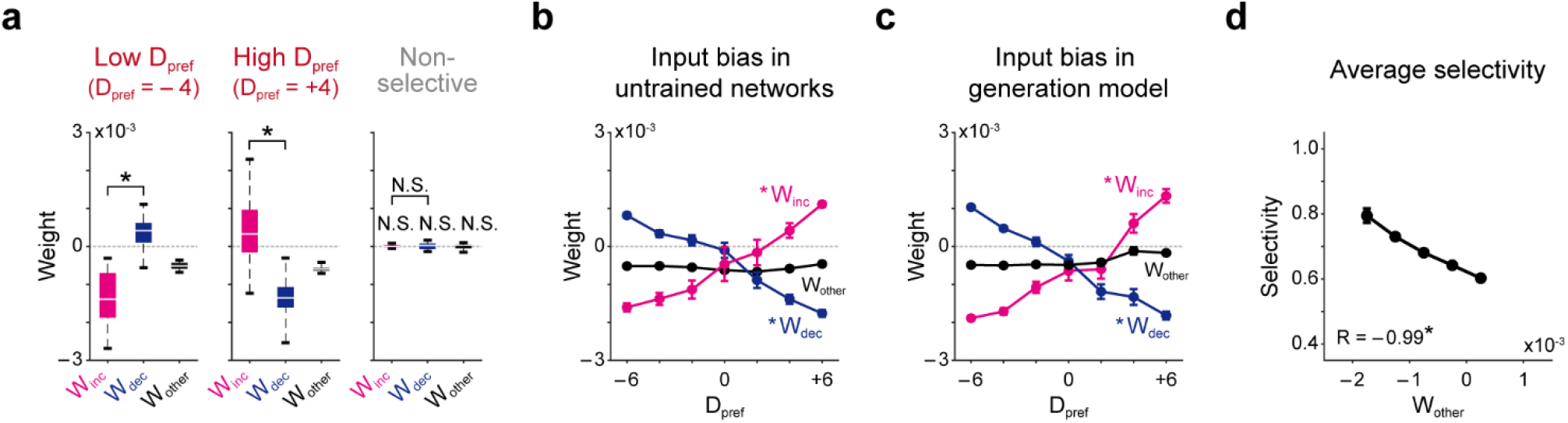
Connectivity weight analysis for difference units. (a) Average weight distribution from Layer 2 units to Layer 3 difference units (D_pref_ = −4, +4). Note that difference units tuned to −4 receive stronger inputs from decreasing units than increasing units, whereas difference units tuned to +4 show the opposite tendency (left, W_inc_ < W_dec_, *P = 1.21 ×10^-8^, middle, W_inc_ > W_dec_, *P = 1.62×10^-8^, paired t-test, N_net_ = 20, N.S., P > 0.05). (b) Average values of W_inc_, W_dec_, and W_other_ of connections from units in Layer 2 to difference units in Layer 3 with different preferred difference values. Similar to Fig. 3f, the difference units with low D_pref_ values receive stronger inputs from decreasing units (blue) and weaker inputs from increasing units (magenta), whereas units with high D_pref_ values receive stronger inputs from increasing units and weaker inputs from decreasing units (*P = 3.97×10^-4^, Mann-Kendall test for increasing/decreasing tendency, N_net_ = 20). (c) Estimated input bias from the generation model. The format is identical to that in **b**. (d) A negative correlation between W_other_ and the average selectivity of the difference units is observed (*P = 6.24×10^-4^, R = −0.99 between the average selectivity and W_other_, Pearson correlation).

**Figure S7.**
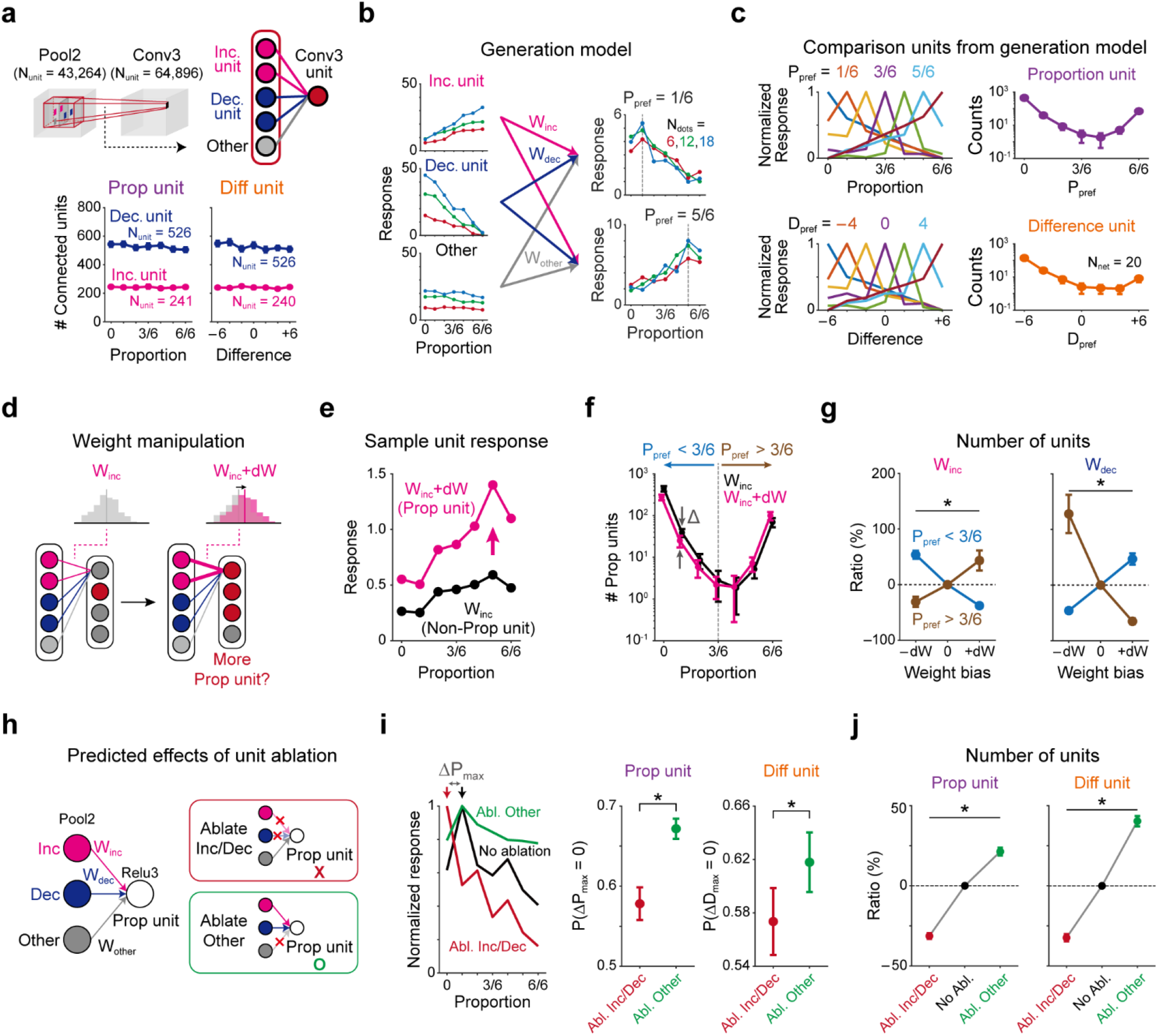
Weight summation model, weight manipulation, and unit ablation test. (a) Top: Illustration of the generation model in a convolutional neural network. Bottom: The number of Pool2 increasing/decreasing units connected to a Relu3 proportion (or, difference) unit was estimated in untrained networks. (b) Generation model. Responses for proportion (difference) stimulus sets are summed and the ReLU function is applied. Selective units were determined using the same method used in the untrained network analysis. (c) Average tuning curves and the distribution of comparison units from the generation model. (d) Weight manipulation scheme. The weights between increasing source units and a target unit are shifted by dW_inc_. (e) Sample unit response of proportion units generated due to the weight manipulation. (f) Change in the number of emerging comparison units due to a positive dW_inc_. Note that more proportion units with high P_pref_ values (P_pref_ > 3/6) were generated than those with low P_pref_ values when the wiring strength from the increasing units (W_inc_) was increased manually. (g) Changes in the number of quantity-comparison units by weight manipulation (*P < 1.86×10^-21^, one-way repeated measure ANOVA with Tukey’s *post-hoc* test). (h) Ablation of increasing/decreasing units and corresponding predictive effects. (i) Sample responses (left) and changes in the peak position after ablation (right). The peak positions of proportion (difference) units were altered significantly more when increasing/decreasing units were ablated, compared to the ablation of non-increasing/decreasing units (*P = 3.41 ×10^-6^ and *P = 0.048 for proportion and difference units, paired t-test, N_net_ = 20). (j) Number of comparison units under different ablation conditions. The emergence probability significantly decreased when all increasing and decreasing units were ablated and significantly increased when all other units were ablated (*P < 3.96×10^-24^, one-way repeated measure ANOVA with Tukey’s *post-hoc* test).

**Figure S8.**
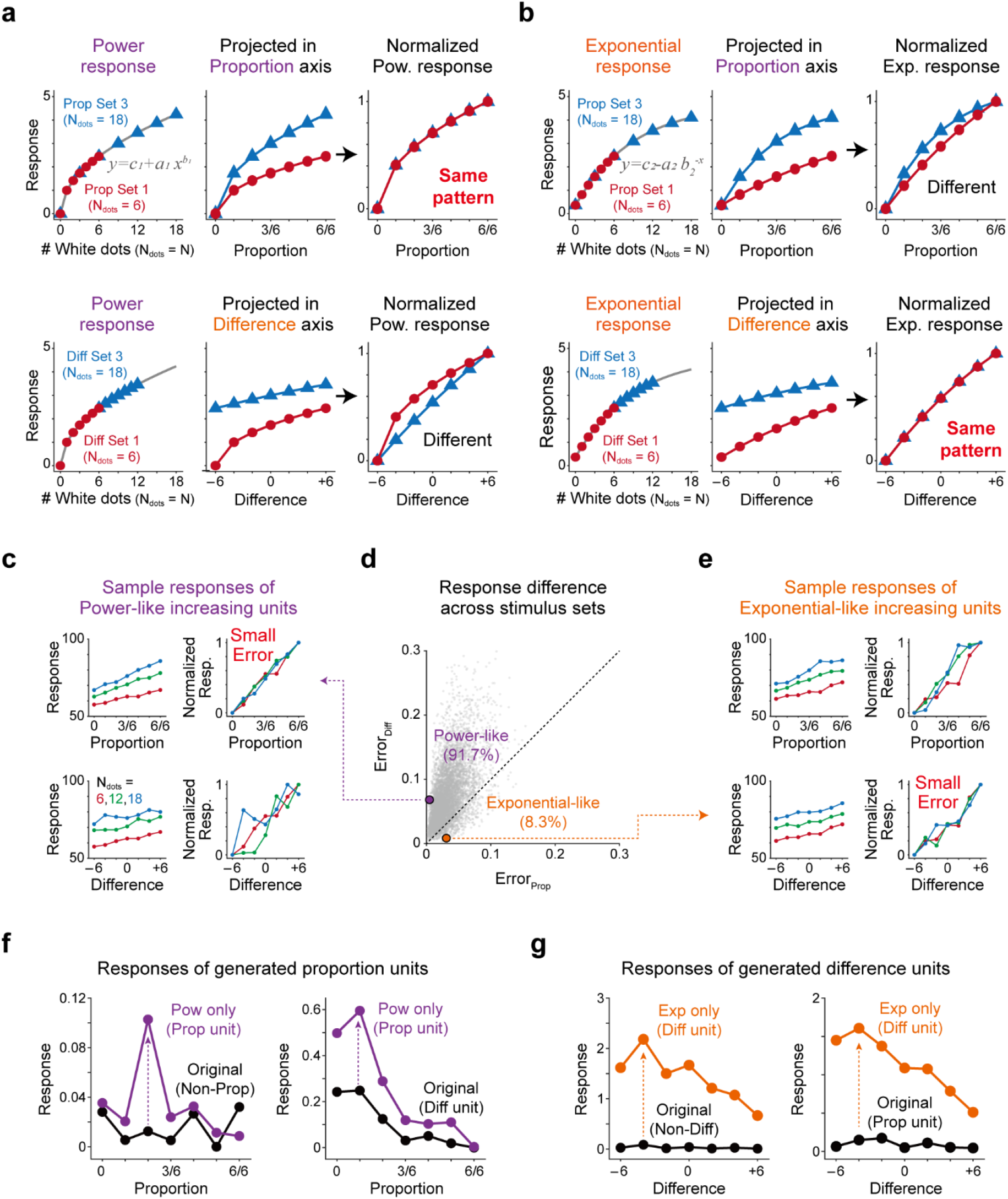
Detailed characteristics of power-like and exponential-like responses generating proportion-selective and difference-selective units. (a) A unit with a response increases by a power function when the number of white dots increases. Proportion stimulus sets with 6 and 18 dots in total are indicated by the red circle and blue triangle, respectively. Response curves projected onto the proportion axis have a similar pattern, becoming identical when normalized (top). In contrast, for difference stimulus sets with 6 and 18 dots in total, the response patterns are different in the two sets and cannot be identical when normalized (bottom). (b) Similar to **a**, but for responses as an exponential function. Exponentially increasing response patterns are different on a proportion-scale axis (top) but become identical on a difference-scale axis (bottom). (c-e) Classification of increasing and decreasing units as power-like and exponential-like units due to the error between normalized responses for proportion and difference stimulus sets. For example, a unit showing a smaller error for a proportion stimulus set than that for a difference stimulus set is considered as a power-like unit. (f-g) Sample responses demonstrate that the type of selective unit can change due to the ablation of power-like or exponential-like units. (f) Responses of proportion units generated by ablation. A non-selective unit (left) or a difference-selective unit (right) becomes a proportion-selective unit when only power-like units are connected. (g) Responses of difference units generated by ablation. A non-selective unit (left) or a proportion-selective unit (right) becomes a difference-selective unit when only exponential-like units are connected.

## Supplementary Notes

### Analytic solution for the emergence of proportion and difference selectivity from the weighted sum of monotonically increasing and decreasing responses

#### 1. Notations

|  |  |
| --- | --- |
| $N$ | Total number of dots in an image ($N \geq 2$) |
| $x$ | Number of white dots in an image ($0 \leq x \leq N$) |
| $f(x), g(x)$ | Input response with respect to the number of white dots ($x$)<br>Both $f(x), g(x)$ are monotonically increasing and concave functions for all $x$. |
| $w_1, w_2$ | Weight connecting input and output units |
| $F(x; N), G(x; N)$ | Response of an output unit that receives weighted input responses<br>$F(x; N) = w_1 f(x) + w_2 f(N - x)$, $G(x; N) = w_1 g(x) + w_2 g(N - x)$<br>where $F$ and $G$ represent the responses of proportion and difference units, respectively |

#### 2. Nonlinear input responses

If a proportion- or difference-selective response can correspondingly be represented by the weighted sum of one increasing response (*f*(*x*) or (*g*(*x*) and one decreasing response (*f*(*N – x*) or *g*(*N – x*)), both responses should be nonlinear functions. Otherwise, *F*(*x; N*) and *G*(*x; N*) become linear functions. Accordingly, both functions cannot have a maximum peak in (0, *N*).

To show this, we can assume that (*f*(*x*) is a linear function. Then, *F′*(*x*) = *w*_1_*f′*(*x*) – *w*_2_*f′*(*N – x*) = *const*, because both *f′*(*x*) and *f′*(*N – x*) are constant for all *x*. Thus, *F(x*) is a linear function as well, implying that the maximum peak occurs either when *x* = 0 *or N*, i.e., a decreasing or increasing function, respectively.

#### 3. Shape of nonlinear responses necessary for proportion and difference selectivity

To find the functional form of a monotonic response that generates proportion selectivity, we found the condition such that *F′*(*x_p_; N*) = 0 *for x_p_* = *pN*, where *p* represents the preferred proportion.

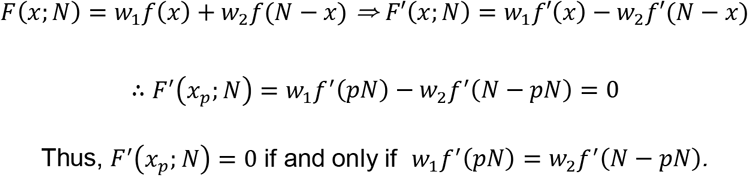

Here, if we assume *f*(*x*) = *x^α^* (0 < *α* < 1), then (LHS) = *w*_1_(*pN*)*^α^* = *w*_1_*p^α^N^α^*, (RHS) = *w*_2_(*N – pN*)*^α^* = *w*_2_(1 – *p*)*^α^N^α^*. Thus, the above equation holds when 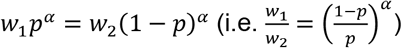.

Hence, a function satisfying *f′*(*x*) = *x^α^* can generate proportion selectivity (e.g., a power/log function).

Similarly, we found a condition, for difference selectivity, such that *G′*(*x_d_; N*) = 0, where *x_d_* – (*N – x_d_*) = *d* and where *d* represents the preferred difference.

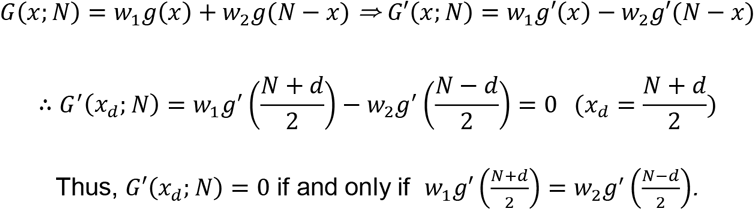

Because *N* and *d* are both constants, we can rewrite *w*_1_*g′*(*t + d*) = *w*_2_*g′*(*t*) for simplicity. Here, if we assume *g′*(*x*) = *e^-x^*, then (LHS) = *w*_1_*e*^-(*t+d*)^ (RHS) = *w*_2_*e^-t^*. Thus, the above equation holds when 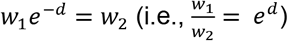.

Hence, a function satisfying *g′*(*x*) = *e^-x^* can generate difference selectivity (e.g., an exponential function).

## Notes

### Competing Interest Statement

The authors have declared no competing interest.

